# Overlapping upstream ORFs repress translation and expand proteome diversity in Arabidopsis

**DOI:** 10.64898/2026.07.14.738309

**Authors:** Hsin-Yen Larry Wu, Yu-Hsuan Cheng, Isaiah D. Kaufman, Qiaoyun Ai, Polly Yingshan Hsu

## Abstract

Upstream open reading frames (uORFs) are widespread cis-regulatory elements that modulate translation initiation of downstream main ORFs (mORFs). Among them, overlapping uORFs (ouORFs) that overlap with mORFs are predicted to exert the strongest translational repression, yet they remain largely unexplored because of the difficulty of their identification. Here, we developed complementary computational approaches to systematically identify translated ouORFs from super-resolution ribosome profiling data in Arabidopsis. We identified 965 translated ouORFs alongside 7,180 canonical non-overlapping uORFs (nuORFs). We found that ouORFs exert substantially stronger translational repression than nuORFs, and that this repression depends primarily on Kozak context rather than uORF length. In addition, genes containing ouORFs or nuORFs have weaker mORF Kozak contexts than genes without uORFs, which may further reduce mORF translation. Moreover, ouORF translation promotes initiation downstream of the annotated mORF start codon, generating N-terminally truncated protein isoforms with altered domain composition and subcellular localization. Using ATPS2 as an example, we demonstrate that ouORF translation regulates alternative translation initiation to control the balance between chloroplast and cytosolic protein isoforms. Together, our findings establish ouORFs as a versatile class of translational regulatory elements that coordinate both protein abundance and protein diversity, providing the first genome-wide characterization of translated ouORFs in plants.

## INTRODUCTION

Translation initiation is a major regulatory step in eukaryotic gene expression and is frequently modulated by upstream open reading frames (uORFs) (1). uORFs are short ORFs that initiate at an AUG or near-cognate codon in the 5′ untranslated region (5′ UTR) upstream of the main ORF (mORF). Their translation generally reduces downstream protein production through several mechanisms, including ribosome drop-off, ribosome stalling, and limited reinitiation (1–3). High-resolution ribosome profiling (Ribo-seq) has enabled the genome-wide identification of thousands of translated uORFs in plants, revealing that they are widespread among genes with important biological functions (4–9). Many uORFs act conditionally, regulating mORF translation in response to metabolites, developmental cues, or environmental signals (1). Their regulatory strength is influenced by several sequence features, including start-codon (Kozak) context and uORF length. Conversely, cells can modulate uORF activity by altering their inclusion in transcripts through alternative transcription start site (TSS) usage or alternative splicing of the 5′ UTR (10, 11). Naturally occurring variants that create, remove, or modify uORFs contribute to phenotypic diversity (12–14), and uORFs have emerged as promising targets for gene engineering and crop improvement (15–17). .

Most uORFs initiate and terminate upstream of the annotated mORF start codon and are referred to as non-overlapping uORFs (nuORFs) (**Fig. 1A**). In contrast, overlapping uORFs (ouORFs) initiate within the 5′ UTR, extend through the mORF start codon, and terminate within the coding sequence (**Fig. 1A**). This architecture fundamentally alters the mechanism of uORF- mediated translational regulation. Following translation of a nuORF, the post-termination 40S ribosomal subunit can remain associated with the mRNA and reinitiate at the downstream mORF. Reinitiation efficiency depends on several factors, including uORF length, intercistronic distance, and the availability of initiator tRNA and initiation factors for reloading onto the 40S subunit (18–20). In contrast, because translation of an ouORF proceeds through the annotated mORF AUG, reinitiation is precluded at the annotated start site. Consequently, ouORFs are predicted to exert stronger translational repression than canonical nuORFs (21). Consistent with their stronger functional impact, ouORFs are under stronger purifying selection in human populations (22).

**Fig. 1.**
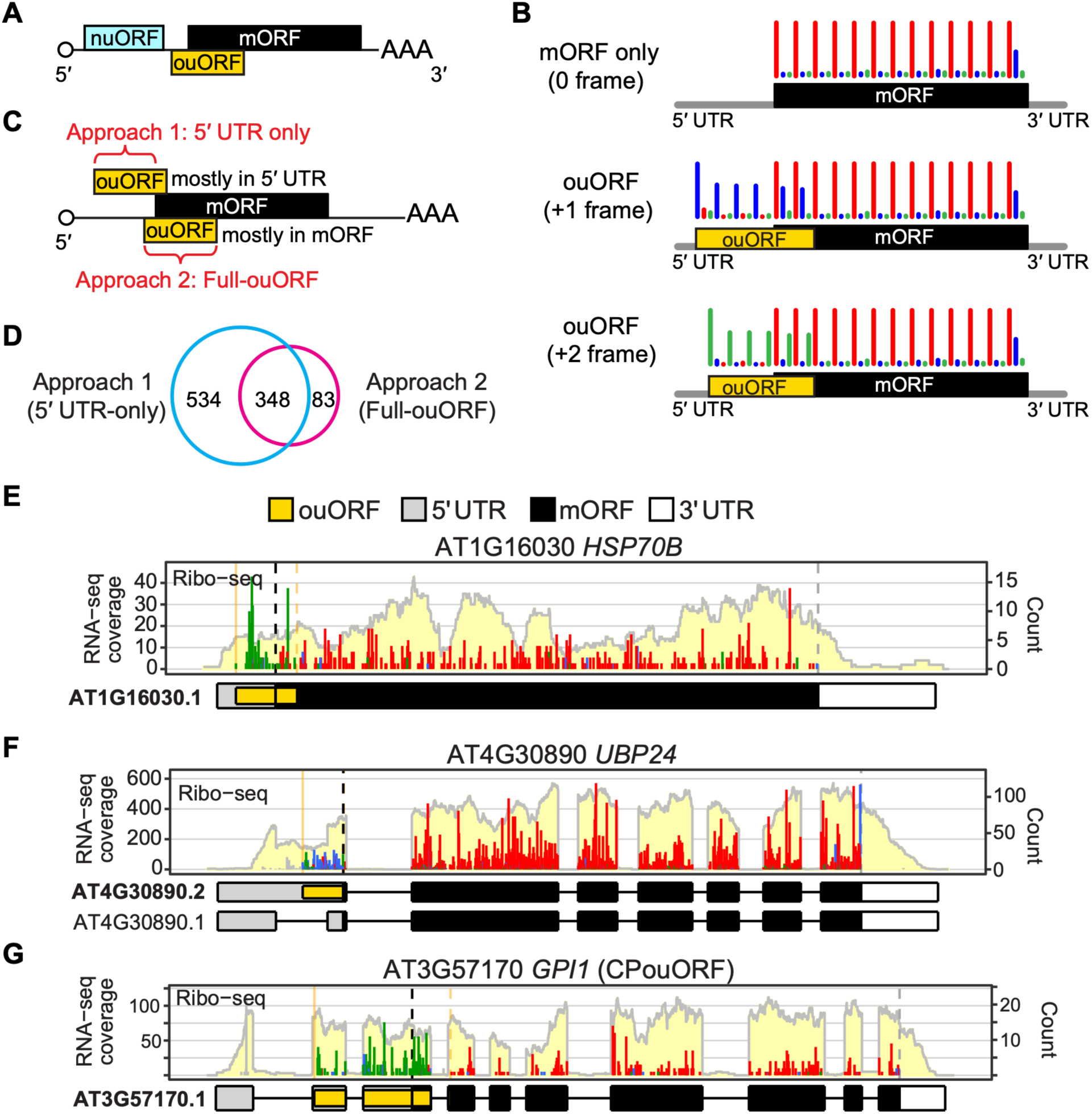
Genome-wide identification of translated ouORFs. (A) Schematic of a nuORF and ouORF. (B) Schematic illustration of expected Ribo-seq profiles for genes translating only the mORF or with a +1- (blue) or +2-frame (green) ouORF. Ribo-seq reads are color-coded by reading frame (red, annotated mORF frame; blue, +1; green, +2). (C) Method 1 scores only the 5′ UTR portion of the ouORF and is likely optimal for ouORFs that span mostly within the 5′ UTR, whereas Method 2 scores the entire ouORF and is likely optimal for ouORFs that span mostly within the mORF. (D) Overlap between ouORFs identified by the two methods. (E–G) Visualization of ouORF examples, including the CPouORF in *GPI1*. *ggRibo* plots (66) show RNA-seq coverage (light yellow background), Ribo-seq reads color-coded by reading frames (relative to the mORF as in Fig. 1B), and gene models with ouORFs highlighted. Black/gray dashed lines mark mORF start/stop codons, and solid/dashed orange lines mark ouORF start/stop codons.

Despite their potential importance, ouORFs have remained largely unexplored in plants. Conventional uORF identification using Ribo-seq relies on the strong 3-nucleotide periodicity of ribosome footprints, which correspond to ribosomes’ codon-by-codon movement within a translated ORF (23, 24). Because ouORFs overlap the mORF in a different reading frame, ribosome footprints from the two ORFs are superimposed, disrupting the apparent periodicity of the ouORF (**Fig. 1B**) and preventing most existing algorithms from detecting them. To our knowledge, only two ouORFs have been functionally characterized in plants. The Arabidopsis *PHOSPHATE1* (*PHO1*) ouORF represses *PHO1* translation and affects shoot phosphate accumulation and growth under phosphate limitation (25). A naturally occurring ouORF in *PETAL LOBE ANTHOCYANIN* (*PELAN*) reduces *PELAN* translation and flower pigmentation in monkeyflower (*Mimulus parishii*) (26). Thus, the prevalence, regulatory properties, and broader biological functions of ouORFs remain largely unknown.

To bridge this gap, here we developed complementary computational approaches and used improved super-resolution Ribo-seq data to systematically identify translated ouORFs in Arabidopsis. This ouORF catalog enabled us to compare the regulatory properties and mechanisms of translational repression between ouORFs and canonical nuORFs. Our results demonstrate that ouORFs not only repress mORF translation more strongly than canonical nuORFs but also promote downstream translation initiation, generating alternative protein isoforms with altered protein domain composition and distinct subcellular localizations.

Together, these findings establish ouORFs as a potent and versatile class of translational regulatory elements.

## RESULTS

### Genome-wide identification of translated ouORFs

Unlike nuORFs that are entirely upstream of the mORF, ouORFs partially overlap mORFs and use the +1 or +2 frame, instead of frame 0 (**Fig. 1A, B**). The overlapping and out-of-frame features disrupt the overall 3-nt periodicity within the ouORFs and prevent their effective identification by conventional uORF identification tools that rely on 3-nt periodicity.

To systematically identify translated ouORFs, we applied two complementary approaches (**Fig. 1C**) and an optimized CiPS pipeline to our recent improved Ribo-seq data in Arabidopsis (9). We reasoned that if an ouORF spans mostly in the 5’ UTR, its Ribo-seq reads in the 5’ UTR alone should exhibit clear 3-nt periodicity that permits identification, so our first approach scores the in-frame information within the 5’ UTR only (**Fig. 1C, top)**. In contrast, if an ouORF spans mostly within the mORF, the mixing signal in two frames will reduce the apparent 3-nt periodicity, and its reads in the 5’ UTR are likely insufficient for identification (**Fig. 1C, bottom)**. To tackle this issue, we developed the second approach, which scores the full ouORF range using adjusted CiPS criteria with more relaxed in-frame cutoff but more stringent length and density requirements (**Fig. S1–S2** for detailed workflow and criteria). We predicted all AUG-initiated ORFs in annotated 5’ UTRs and scored each candidate using the two approaches. Candidates with a stop codon upstream of the main AUG were classified as canonical nuORFs; those whose codons continue past the main AUG and terminate within the mORF were classified as ouORFs (**Fig. 1A**).

In total, we identified 965 translated ouORFs in 919 genes (**Table S1**). The two ouORF identification approaches were complementary: 534 ouORFs were identified by the ‘5’-UTR-only’ approach alone, 83 by the ‘full-ouORF’ approach alone, and 348 by both approaches (**Fig. 1D**). We also identified 7,180 translated nuORFs in 4,810 genes using the first approach (**Table S2**). Visualizing the Ribo-seq/RNA-seq profiles of these ouORFs using *ggRibo* confirmed their active translation in either +1 or +2 reading frames relative to the mORF (**Fig. 1E–G** and other *ggRibo* plots throughout the article, e.g., **3A–C**; **5D–F**). The profiles also revealed examples of ouORFs that mostly span the 5’ UTR (e.g., **Fig. 1F, G**) or mORF (e.g., **Fig. 3A, 5E**).

**Fig. 2.**
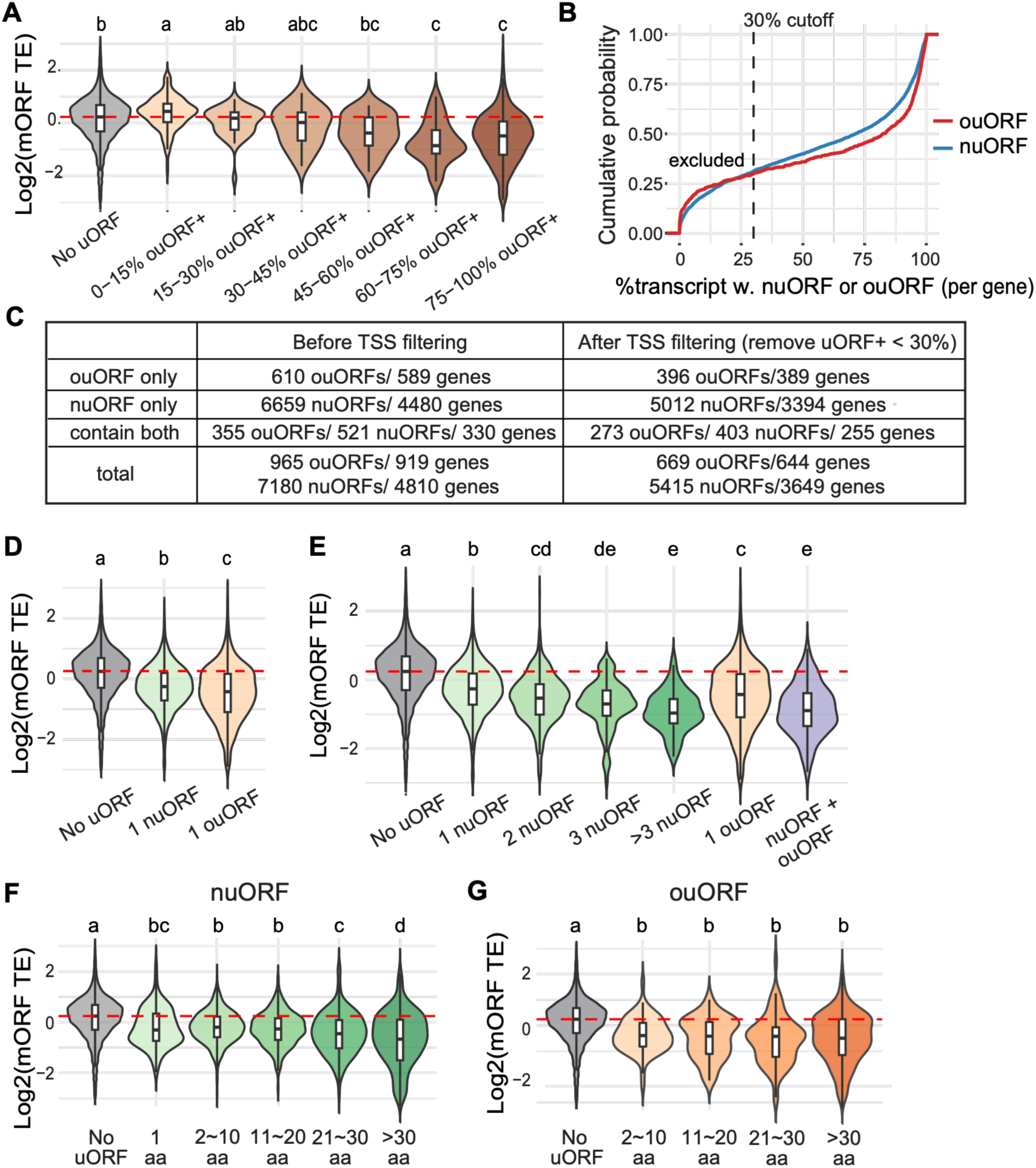
TSS filtering reveals that ouORFs are associated with stronger, length- independent translational repression than nuORFs. (A) TSS-mediated ouORF inclusion is tightly associated with translational repression. Distribution of mORF translation efficiency (TE) for genes lacking uORFs and genes with different percentages of ouORF-containing (ouORF+) transcripts, determined by the proportion of CAGE signal upstream of the ouORF start codon. (B) Cumulative distribution of the percentage of uORF-containing transcripts (nuORF+ or ouORF+) per gene, determined by the proportion of CAGE signal upstream of the uORF start codon. The dashed line indicates the 30% cutoff; 31.2% of nuORF genes and 30.1% of ouORF genes fell below this threshold and were excluded from downstream analyses. (C) Numbers of nuORF- and ouORF-containing genes before and after TSS filtering. (D–G) Distribution of mORF TE after TSS filtering, comparing genes lacking uORFs with genes containing a single nuORF or ouORF (D), genes with different numbers and classes of uORFs (E), and genes containing a single nuORF (F) or ouORF (G) grouped by uORF length. The red dashed line indicates the median TE of genes lacking uORFs. Different letters indicate statistically significant differences (ANOVA followed by Tukey’s HSD test, p < 0.05).

**Fig. 3.**
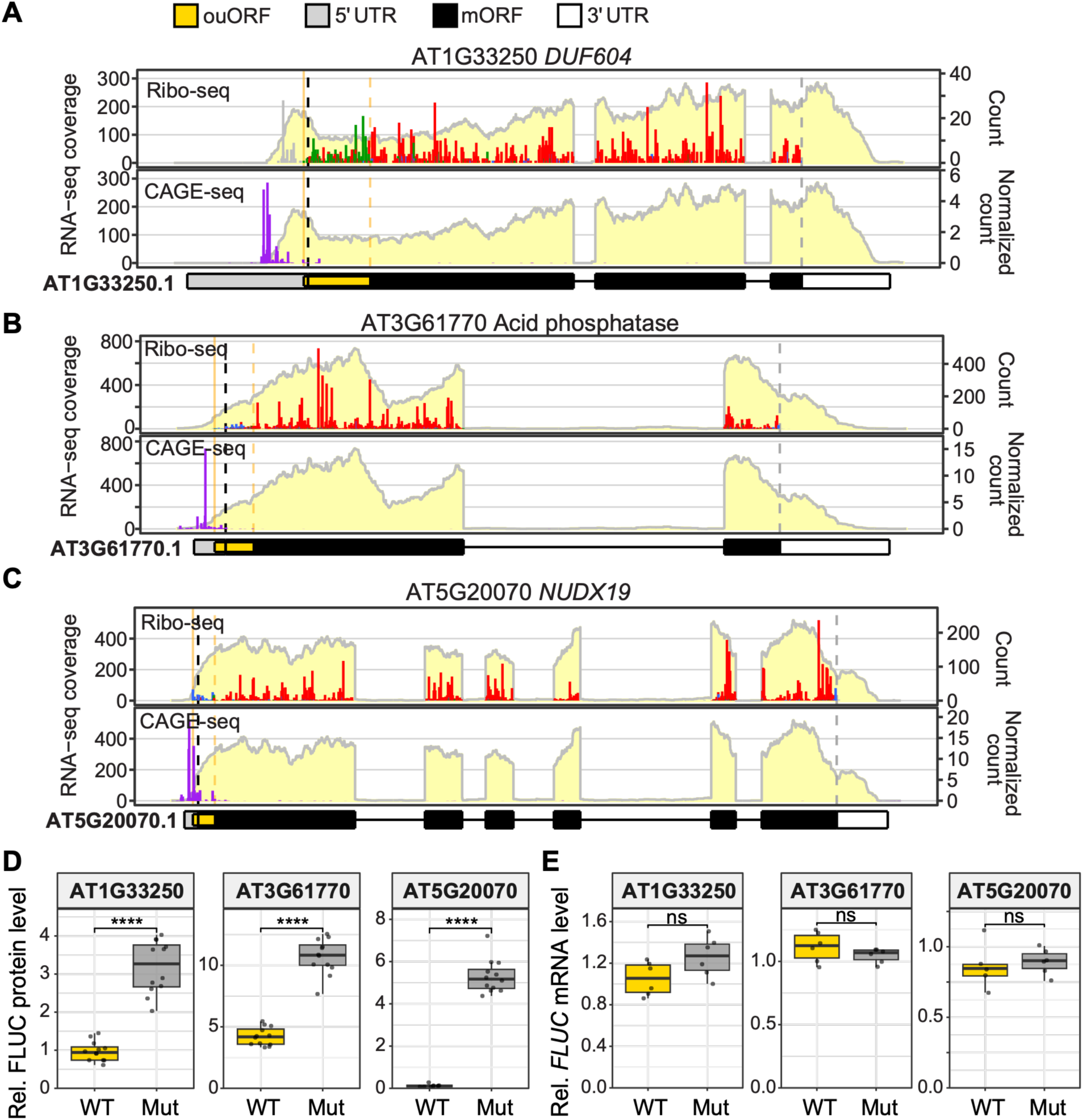
Validation of ouORF-mediated repression of downstream mORF translation. (A–C) Expression profiles of three ouORF-containing genes visualized using *ggRibo*. Data presentation is described in the Fig. 1E legend. (D) Dual-luciferase assay comparing 5′ UTRs containing the wild-type (WT) ouORF or a mutated ouORF (Mut; start codon ATG>AGG). Mutation of the ouORF start codon increases downstream translation. (E) RT-qPCR shows no corresponding change in reporter mRNA levels (ns), indicating that the increase in FLUC protein levels results from translational derepression rather than changes in mRNA abundance. Welch’s two-sample t-test; ****, p < 0.0001; ns, p > 0.05.

Previous work predicted that three AUG-initiated ouORFs encode conserved peptides (conserved peptide ouORFs, or CPouORFs) across plant species (27), but whether these ouORFs are translated in planta has not been confirmed. Our analysis identified all three predicted CPouORFs, from *GPI1* (AT3G57170), a copper amine oxidase (AT2G42490), and *PDBG4* (AT1G11820), and confirmed their active translation in our Ribo-seq data (**Fig. 1G and S3A–C**, Ribo-seq panels). Notably, the *GPI1* CPouORF is translated at a level comparable to that of its annotated mORF (**Fig. 1G**), and its 145-aa peptide is nearly completely conserved across diverse plant species (27), implying this CPouORF peptide may function as an independent protein, similar to the bicistronic *CDC26–TTM3* pair (28) and the *AtMYB51* uORF, which encodes a signaling peptide that triggers systemic stomatal immunity (29).

By overcoming the limitations of conventional uORF identification methods, our integrated approach reveals the hidden landscape of translated ouORFs and provides a foundation for systematically examining their sequence features, translation regulation, and biological functions.

### Isolating ouORFs with CAGE-seq supported transcription start sites

Our recent work revealed that the inclusion of uORFs in mRNAs and their repression are strongly affected by alternative transcription start sites (TSSs), and the dispersed TSSs can mask length-dependent translational repression of uORFs (30). Visual examination of ouORF profiles confirmed that some ouORF genes are similarly influenced by alternative TSSs, as many CAGE-seq reads, which map TSSs, were detected downstream of the ouORF start codon and RNA-seq coverage was markedly reduced near or upstream of the ouORF start codon (**Fig. S4**), indicating that a substantial fraction of transcripts lack the ouORF altogether.

To isolate ouORFs that are frequently retained on mRNAs and therefore have the opportunity to function, we quantified the fraction of transcripts containing each uORF for a given gene using CAGE-seq data (described as %ouORF+, %nuORF+, or collectively %uORF+) (**Tables S1 and S2**) and examined its relationship with mORF translation efficiency (TE). Among genes containing a single uORF, notably, uORFs present on fewer than 30% of transcripts were not associated with a significant reduction in mORF TE (**Figs. 2A and S5**). Above this threshold, mORF TE decreased progressively with increasing %uORF+ for both nuORFs and ouORFs (**Figs. 2A and S5**). Based on the 30% uORF+ threshold, we excluded approximately 31% of nuORF-containing genes and 30% of ouORF-containing genes for subsequent analyses (**Fig. 2B**). This filtering yielded a high-confidence dataset comprising 5,415 nuORFs (3,649 genes) and 669 ouORFs (644 genes) (255 genes containing both classes) (**Fig. 2C**).

Importantly, all three previously predicted conserved peptide ouORFs (CPouORFs) are preferentially retained on mRNAs and show 96–98% ouORF+ (**Fig. S3A–C**, CAGE panels). This supports the idea that functional uORFs are predominantly retained in mRNAs to carry out their biological functions.

### ouORF-containing genes are enriched for protein kinases and phosphatases

We next investigated which biological functions were enriched among our uORF- containing genes using Gene Ontology over-representation analysis. Both nuORF- and ouORF- containing genes were enriched for signaling-related functions (**Table S3**). The strongest shared enrichment was in reversible protein phosphorylation: both gene sets were enriched for protein kinase activity, including serine/threonine kinase activity, as well as for protein serine/threonine phosphatase activity. These results indicate that reversible phosphorylation is a major shared regulatory process associated with both uORF classes.

The two gene sets also differed (**Table S3**). The larger nuORF set exhibited broader functional enrichment, including transcription factor activity (sequence-specific DNA binding and transcription regulator activity), signaling receptors, and signal transduction proteins. Thus, nuORFs are associated with multiple layers of plant regulatory networks, including kinases, phosphatases, receptors, and transcription factors. In contrast, the smaller ouORF set showed the strongest enrichment for protein kinase and phosphatase activities, and notably, poly(A)- specific ribonuclease (deadenylase) activity, which was not enriched among nuORF-containing genes. Genes carrying both nuORF and ouORF were uniquely enriched for phosphatidylinositol (lipid) kinase activity. Collectively, these results suggest that although both uORF classes preferentially regulate signaling genes, ouORFs are more specifically associated with enzymes that control reversible phosphorylation.

### ouORFs are stronger translational repressors

We asked whether ouORFs exert stronger translational repression than nuORFs. We compared genes carrying a single uORF of either class with genes lacking both classes of uORFs (controls) by analyzing mORF TE. Although both uORF classes were associated with reduced mORF TE, significantly stronger repression was observed in ouORFs (**Fig. 2D**). A single ouORF was associated with an mORF TE reduction comparable to that of genes carrying two nuORFs (**Fig. 2E**). The stronger repressive effect of ouORFs is likely due to translation extending past the annotated start codon, thereby precluding reinitiation at the annotated AUG, although reinitiation at downstream AUGs remains possible.

To validate the repressive effect of ouORFs on downstream translation, we performed dual-luciferase assays on three ouORF-containing genes: AT1G33250 (*DUF604*), AT3G61770 (acid phosphatase), and AT5G20070 (*NUDX19*) (**Fig. 3A–C**). For each gene, we inserted the 5′ UTR containing the ouORF and extending through the first four mORF codons upstream of the *FLUC* reporter. We compared the wild-type construct with a mutated version in which the ouORF start codon was disrupted (ATG to A**G**G). Mutating the ouORF significantly increased FLUC activity for all three genes (**Fig. 3D**) without affecting *FLUC* mRNA abundance (**Fig. 3E**), confirming that these ouORFs repress downstream translation.

### OuORF activity is Kozak context-dependent but not length-dependent

We next investigated whether other features influence ouORF-mediated repression. First, we examined the effect of ouORF length on translational repression. Whereas longer nuORFs were associated with progressively stronger repression (**Fig. 2F**) (Wu and Hsu, 2024), ouORF repressiveness showed little dependence on length (**Fig. 2G**). This difference likely reflects distinct mechanisms of repression: because ouORF translation extends through the annotated main AUG, reinitiation at that site is precluded regardless of ouORF length, making repression less dependent on the distance translated before termination than for nuORFs.

Second, we examined the Kozak context of these uORFs. Sequence logos spanning positions −4 to +6 showed that the annotated mORF AUGs of control genes display the canonical optimal Kozak context, characterized by a purine at −3 and a G at +4 (31), whereas nuORF and ouORF start codons exhibited little sequence conservation surrounding the AUG (**Fig. 4A, S6**). Minimal nuORFs (AUG–STOP) were analyzed separately because their Kozak contexts are constrained by the immediate stop codon. Kozak relative strength score further showed that all uORF classes had significantly weaker Kozak contexts than control mORFs, with minimal nuORFs showing the weakest contexts, followed by ouORFs and longer nuORFs (**Fig. 4B**). Among genes containing a single uORF, stronger uORF Kozak contexts were associated with lower mORF TE for both nuORFs and ouORFs (**Fig. 4D, E**), consistent with more efficient uORF initiation leading to stronger repression of downstream translation. Notably, the negative association between uORF Kozak strength and mORF TE was more pronounced for ouORFs than for nuORFs (**Fig. 4D, E**).

**Fig. 4.**
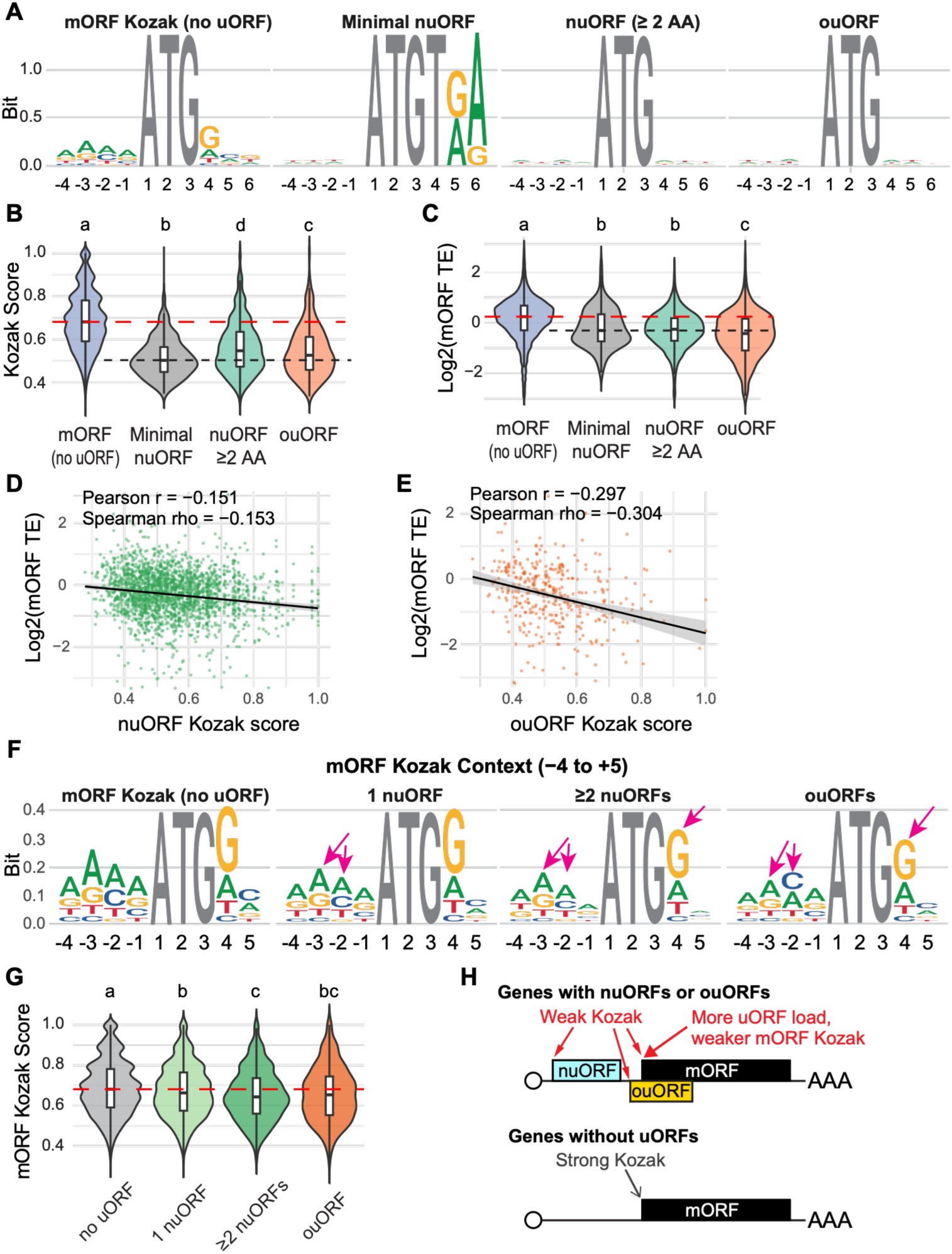
uORF-bearing genes exhibit weaker mORF Kozak contexts. (A) Sequence logos surrounding the start codons of mORFs (uORF-less), minimal nuORFs, nuORFs ≥2 aa, and ouORFs, showing that uORFs generally have weaker Kozak contexts than mORFs. Letter height (bits) indicates information content at each position. (B, C) Kozak score distributions (B) and mORF translation efficiency (TE; C) for the same groups. (D, E) Relationship between uORF Kozak score and mORF TE for genes containing a single nuORF (D) or ouORF (E). Stronger uORF Kozak contexts are associated with lower mORF TE, with a stronger correlation for ouORFs. Black lines represent fitted trends with 95% confidence intervals. (F, G) Sequence logos (F) and Kozak score distributions (G) of mORF start codons from uORF-less genes (control) and genes containing one nuORF, ≥2 nuORFs, or ouORFs. Magenta arrows indicate positions substantially different from the control. (H) Model summarizing the combined effects: uORF-bearing genes have weaker mORF Kozak contexts, with progressively weaker contexts as uORF number increases. Different letters indicate statistically significant differences (ANOVA followed by Tukey’s HSD test, p < 0.05). Red and black dashed lines indicate the median values for mORFs from uORF-less genes and minimal nuORFs, respectively.

Together, these results indicate that ouORF-mediated repression depends more strongly on start-codon recognition than nuORF-mediated repression but is largely independent of uORF length, consistent with a mechanism in which translation extending through the annotated main AUG precludes reinitiation at that site.

### ouORF- and nuORF-bearing genes have weaker mORF Kozak contexts

Less intuitively, previous bioinformatics work reported that the presence of upstream AUGs also correlates with weaker mORF Kozak contexts across diverse eukaryotes (32). We therefore examined the Kozak context of annotated mORF start codons in genes containing nuORFs or ouORFs and found that they were consistently weaker than those of control genes lacking uORFs (**Fig. 4F, G**). Moreover, the mORF Kozak context was progressively weaker in genes containing multiple nuORFs than in those containing a single nuORF (**Fig. 4G, H**). Thus, in addition to repression mediated by uORF translation, a suboptimal mORF Kozak context also contributes to reduced mORF translation.

### ouORF-mediated downstream translation initiation generates N-terminally truncated protein isoforms with altered domain composition and subcellular localization

Although ouORFs preclude reinitiation at the annotated mORF AUG, ribosomes may instead reinitiate at a downstream AUG. When this downstream AUG is in frame with the mORF, translation would produce an N-terminally truncated protein isoform, which can cause loss in protein domains or changes in subcellular localization (**Fig. 5A**). Among 669 translated ouORF-containing genes, 652 contain a downstream in-frame AUG, potentially resulting in N-terminal truncations of 88 aa on average (median 74 aa) (**Table S4**).

**Fig. 5.**
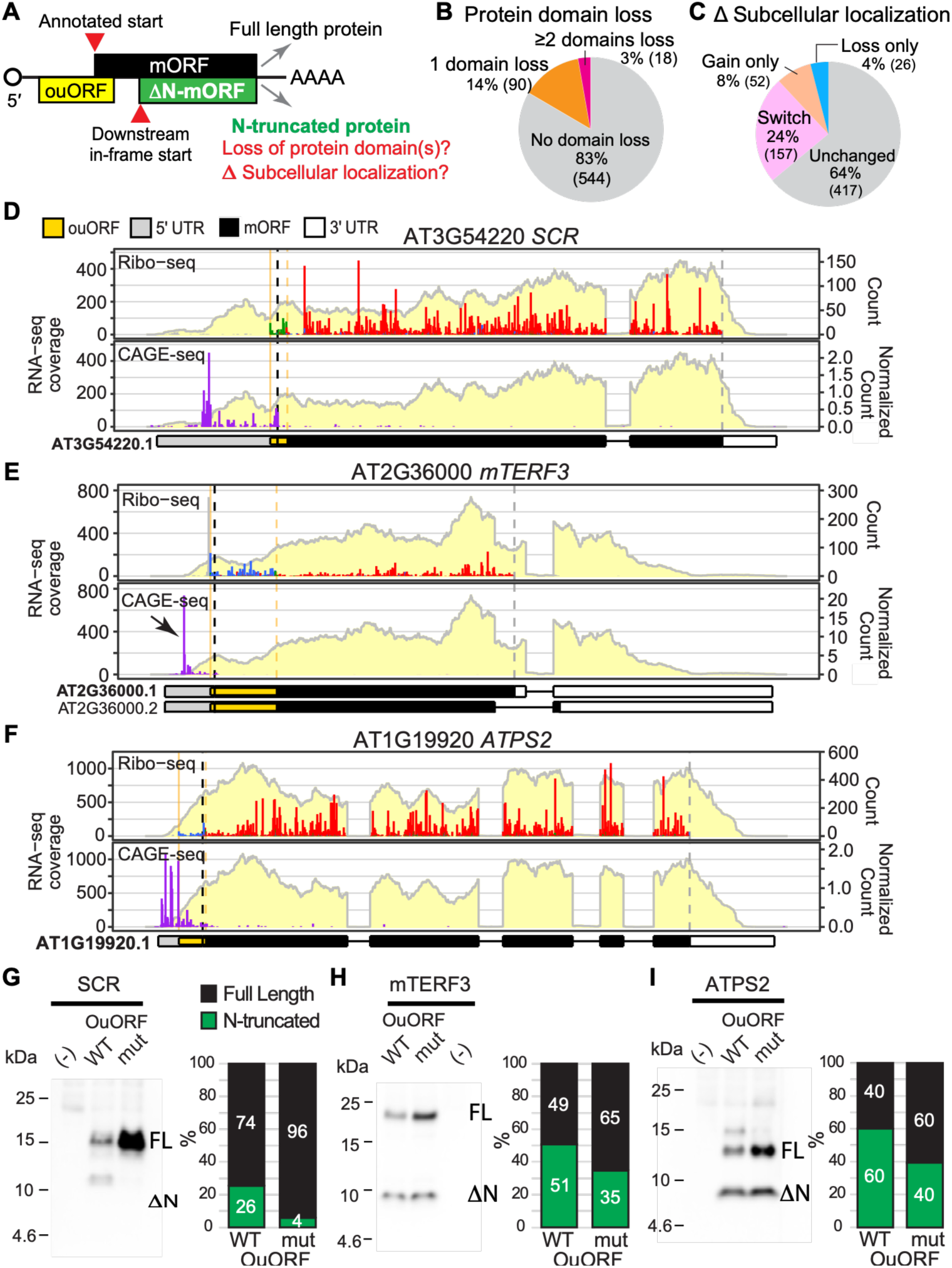
ouORFs promote reinitiation at downstream start codons and generate N-terminal truncated protein isoforms. (A) Schematic illustrating how an ouORF alters mORF start-codon selection. Following ouORF translation, the annotated start codon is bypassed, and translation reinitiates at a downstream AUG, producing an N-terminally truncated protein that may lose functional domains or alter subcellular localization. (B–C) Predicted protein domain loss (B) or change in subcellular localization (C) resulting from ouORF-mediated N-terminal truncation. (D–F) Expression profiles of *SCR*, *mTERF3*, and *ATPS2*, visualized using *ggRibo*. Data presentation is described in the Fig. 1E legend. (G–I) Immunoblots and quantification showing the ratio of full-length to N-terminally truncated protein isoforms for SCR, mTERF3, and ATPS2. Mutation of the ouORF reduces the proportion of the N-terminal truncated isoform across all three genes. Proteins were synthesized by in vitro translation of capped and polyadenylated, in vitro-transcribed mRNAs containing the wild-type ouORF, mutated ouORF (ATG>AGG), or a no-mRNA control (–) using wheat germ extract.

These N-terminal truncations are predicted to cause 17% of ouORF genes to lose one (14%) or ≥2 protein domains (3%) (**Fig. 5B**). The most frequently lost domain, LRRNT_2, is characteristic of LRR-containing receptor-like kinases or receptor proteins, where it mediates protein–protein and ligand interactions. Other frequently lost domains include F-box, transcription factor DNA-binding (e.g., TCP, MYB, MADS-box/SRF-TF, C2H2 zinc finger, BED, and NAM), organellar single-stranded DNA-binding (OSB_SSB), PP2C phosphatase, and protein kinase domains (**Table S4**).

The ouORF-mediated N-terminal truncation may also remove or expose a signal peptide or localization signal, causing the shorter protein isoforms to localize differently from the full-length protein. We examined whether the predicted subcellular localization may change between the full-length and N-truncated protein isoforms for the 652 ouORF genes using DeepLoc2.1 (33). Strikingly, we found that 36% (235/652) had altered predicted localization, with 24% switching localizations, 8% gaining and 4% losing specific subcellular localization (**Fig. 5C**). The most common changes involve redistribution between plastids, mitochondria, or the plasma membrane and the cytosol (**Table S5**).

To test whether ouORFs promote the production of N-terminal truncated proteins, we examined three ouORF-containing genes: AT3G54220 (*SCARECROW*; *SCR*), AT2G36000 (*MITOCHONDRIAL TRANSCRIPTION TERMINATION FACTOR 3*; *mTERF3*), and AT1G19920 (*ATP SULFURYLASE 2*; *ATPS2*) (**Fig. 5D–F**). For each gene, we generated capped and polyadenylated mRNAs containing either the wild-type or a mutated ouORF (ATG to AGG) in the 5′ UTR, together with the first 93–171 codons of the mORF fused to an HA tag. The transcripts were translated in vitro, and protein products were detected by immunoblotting. Consistent with our hypothesis, wild-type constructs produced two protein isoforms corresponding to initiation at the canonical AUG (full-length) and the downstream in-frame AUG (N-terminally truncated). Disrupting ouORF initiation increased the relative abundance of the full-length protein while decreasing the N-terminally truncated isoform (**Fig. 5G–I**), indicating that ouORF translation promotes initiation at downstream in-frame AUG codons. Moreover, the increase in total protein abundance following ouORF mutation provides additional evidence that ouORFs repress overall mORF translation.

Together, these findings indicate that ouORFs not only repress translation at the canonical start codon but also promote downstream AUG usage, expanding protein diversity through alternative N-terminal isoforms.

### OuORF-mediated changes in ATPS2 subcellular localization

A previous study reported that ATPS2, a key enzyme in the sulfate assimilation pathway, localizes to both chloroplasts and the cytosol. The authors proposed that the cytosolic isoform is generated by leaky scanning at the annotated start codon (M1), allowing translation to initiate at downstream in-frame AUGs (M52 or M58) and thereby bypass the N-terminal chloroplast transit peptide (34). Our immunoblotting results (**Fig. 5I**) support an alternative model in which the ouORF promotes downstream initiation by preventing reinitiation at M1. Consistent with this model, the Ribo-seq profile shows that ribosome footprints are more abundant at M52 and M58, whereas initiation at M1 is greatly reduced following ouORF translation, indicating that ATPS2 preferentially initiates at the downstream AUGs (**Figs. 6A** and **5F**).

**Fig. 6.**
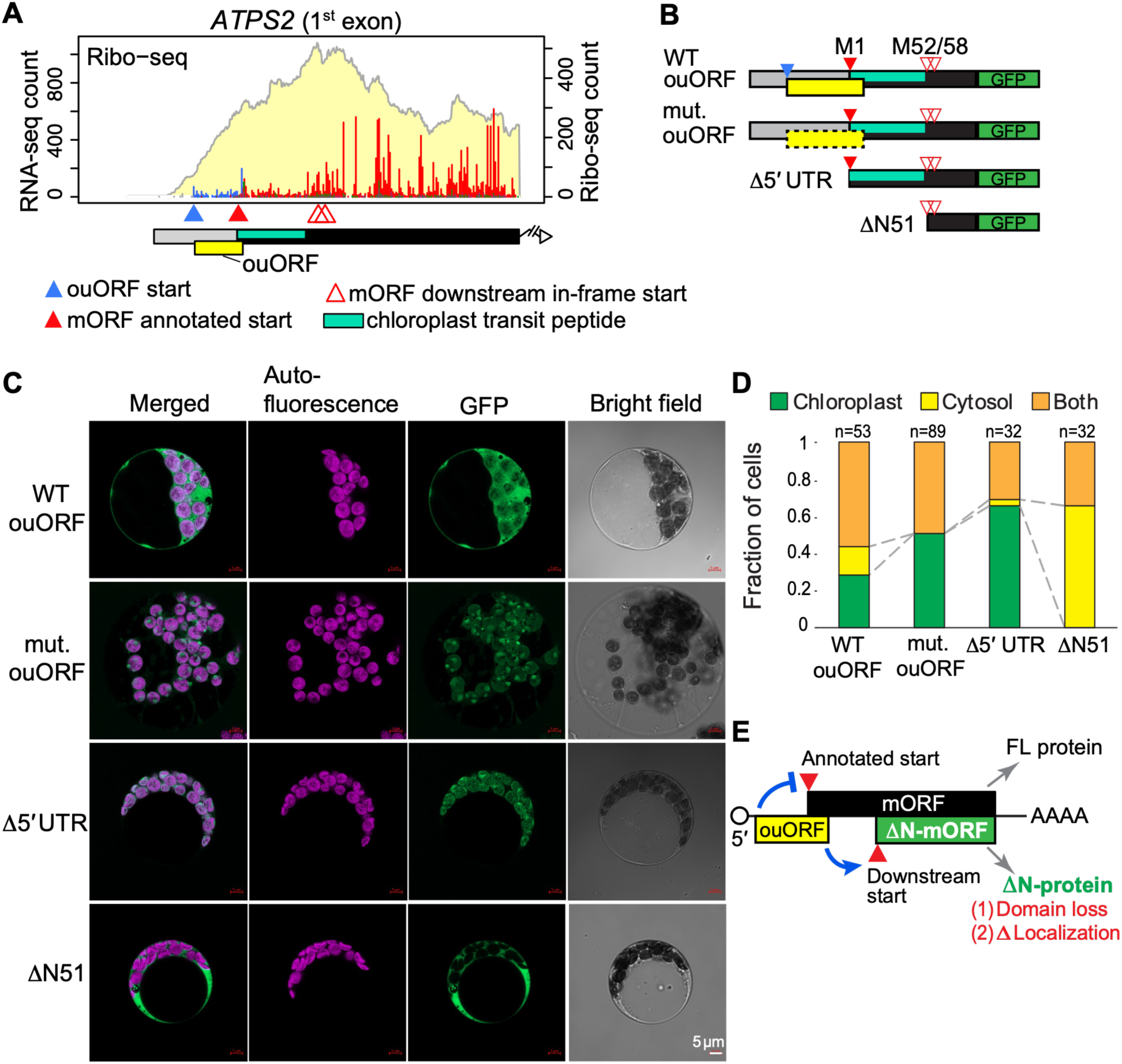
ouORF-mediated downstream translation initiation generates an N-terminally truncated, cytosolic ATPS2 isoform. (A) Enlarged view of the first exon of ATPS2, highlighting the positions of the ouORF, the annotated and downstream mORF start codons, and the chloroplast transit peptide. The same RNA-seq and Ribo-seq data shown in Fig. 5D are displayed. (B) Schematic of the ATPS2-GFP constructs used for localization assays: WT ouORF, mutated ouORF (start codon ATG>AGG), Δ5′ UTR, and ΔN-terminal. Red triangles indicate AUG codons, and green boxes represent GFP. (C) Representative confocal images of Arabidopsis leaf protoplasts expressing each construct, showing merged images, chlorophyll autofluorescence, GFP fluorescence, and bright-field images. (D) Quantification of GFP localization patterns shown in E. Cells were classified as chloroplast, cytosol, or both. Numbers above the bars indicate the number of cells analyzed. (E) Proposed model of ouORF-mediated repression of canonical mORF initiation and indirect promotion of reinitiation at a downstream start codon, generating N-terminally truncated proteins that may lose protein domains and/or exhibit altered subcellular localization.

To determine whether ouORF-mediated start-site selection alters ATPS2 subcellular localization, we expressed four GFP reporter constructs in Arabidopsis leaf protoplasts: wild-type ouORF or mutated ouORF (start codon ATG to TAG) in the 5′ UTR, a 5′ UTR deletion (Δ5′ UTR), and an N-terminal deletion (ΔN) (**Fig. 6B**). Consistent with the previous study (34), in the wild-type construct, ATPS2 is localized to both chloroplasts and the cytosol (**Fig. 6C, D**). In contrast, mutating the ouORF increased chloroplast localization while reducing cytosolic localization. Deletion of the 5′ UTR resulted predominantly in chloroplast localization, whereas deletion of the N-terminal transit peptide produced predominantly cytosolic localization (**Fig. 6C, D; Fig. S7**). Together, these findings demonstrate that ouORF-mediated start-site selection controls the balance between chloroplast and cytosolic ATPS2 isoforms.

## DISCUSSION

Our study provides the first systematic characterization of translated ouORFs in plants and establishes them as potent translational regulatory elements that regulate both protein abundance and protein diversity. By integrating new identification approaches, TSS analysis, and improved super-resolution Ribo-seq, we establish a framework for investigating their regulation and biological functions.

Consistent with expectations, we found that ouORFs exert stronger translational repression than nuORFs, in agreement with studies in vertebrates showing that ouORFs are among the strongest predictors of translational repression and are subject to stronger purifying selection (21, 22). Our analyses revealed that ouORFs repress translation through a mechanism fundamentally distinct from that of canonical nuORFs. Whereas nuORF-mediated repression depends strongly on uORF length because longer uORFs reduce the probability of efficient reinitiation (19), ouORFs exhibited little length dependence. Instead, ouORF activity was determined primarily by start-codon context, consistent with a model in which initiation at an ouORF commits the ribosome to translate through the annotated mORF AUG, thereby preventing initiation at that site. We further found that genes containing either nuORFs or ouORFs possess weaker mORF Kozak contexts than genes lacking uORFs, suggesting that inefficient initiation at the annotated start codon provides an additional, uORF-independent layer of translational regulation.

Random fluctuations in mRNA abundance are a major source of gene-expression noise. Reduced protein production per mRNA constrains translational burst magnitude and dampens the transmission of this mRNA-level noise to protein abundance (35–37). Accordingly, uORF-mediated translational repression can reduce protein-expression noise and stabilize protein abundance, consistent with broader evidence that uORFs buffer variation in downstream translation (38, 39). Our finding that uORF-containing genes also have weaker mORF Kozak contexts points to an additional layer of translational repression that could reinforce this buffering effect. The preferential enrichment of uORFs, particularly ouORFs, in protein kinases and phosphatases further suggests that these regulatory mechanisms are concentrated within signaling networks, where precise control of protein abundance is likely especially important.

Beyond regulating protein abundance, our results reveal that ouORFs expand proteome diversity by promoting the production of N-terminally truncated protein isoforms. Alternative N termini are an important source of protein diversity and can alter protein stability, interaction partners, regulatory activity, and subcellular localization (40, 41) . Translation initiation site sequencing (TI-seq) and N-terminal proteomics have shown that alternative N-terminal proteoforms are widespread in plants (42, 43). Although these proteoforms have traditionally been attributed to mechanisms such as leaky scanning, alternative TSS usage, or alternative splicing (34, 44–48), our findings identify ouORF translation as an additional mechanism that promotes downstream translation initiation and generates alternative protein isoforms.

More broadly, ouORF-mediated downstream translation initiation may represent a widespread mechanism for expanding protein localization and functional diversity in plants. Protein function is often closely linked to subcellular localization, and the same protein can perform distinct molecular functions in different cellular compartments. For example, plastid-localized WHIRLY1 functions in plastid nucleoid organization and chloroplast development, whereas nuclear WHIRLY1 regulates salicylic acid homeostasis and leaf senescence through transcriptional control (49, 50). Likewise, ouORF-mediated domain loss may generate protein isoforms resembling microproteins, which are typically single-domain proteins that act as dominant-negative regulators by sequestering their multidomain homologs (51). The LITTLE ZIPPER proteins exemplify this mechanism: they retain the leucine zipper domain required for interaction with HD-ZIPIII transcription factors but lack the homeodomain, thereby inhibiting HD-ZIPIII activity through nonfunctional heterodimer formation (52, 53). It is possible that ouORF- mediated loss of transcription factor domains could generate dominant-negative isoforms with analogous regulatory functions. Together, these findings suggest that ouORF-mediated downstream translation initiation represents a previously unrecognized mechanism for expanding proteome diversity by generating alternative protein isoforms with distinct localization and altered biological functions.

In summary, ouORFs constitute a distinct class of translational regulatory elements that regulate both protein abundance and protein diversity by redirecting translation to downstream start codons, thereby generating alternative protein isoforms with altered domain composition, subcellular localization, and functional complexity (**Fig. 6E**).

## MATERIALS AND METHODS

### Genome annotation

Transcript models, coding sequences (CDSs), exons, and 5′ UTRs were retrieved from the Araport11 annotation (54).

### Identification of translated ouORFs and nuORFs

Improved super-resolution Ribo-seq and accompanying RNA-seq data from Arabidopsis Col-0 seedlings were previously described (9) (GEO accession GSE183264). All read preprocessing, including trimming, alignment, and P-site assignment, was performed previously, and the processed P-site table was used directly in this study.

Two key improvements to the CiPS method (9) were introduced. First, all annotated transcript isoforms from Araport11 were used instead of the simplified GTF annotation employed previously. Second, nuORF stop codons were allowed to overlap the annotated mORF AUG (**Fig. S2B**).

Candidate AUG-initiated uORFs were identified by scanning annotated 5′ UTRs, extended into the mORF, using ORFik (55). CiPS scores each candidate ORF for active translation using three metrics—in-frame count, in-frame percentage, and in-frame site percentage—calculated with a vectorized counting engine that maps each P-site to a codon position within the ORF and classifies it as in-frame or out-of-frame, except that those mapping to the second nucleotide of the terminal sense codon are counted as in-frame (9). Candidates terminating upstream of the annotated mORF AUG were classified as nuORFs, whereas those extending through the annotated AUG and terminating within the mORF were classified as ouORFs. Because ribosome footprints from ouORFs are split between the 5′ UTR and the mORF-overlapping region, ouORFs were evaluated using two complementary approaches: a 5′- UTR-only approach (criteria: in-frame count ≥10, in-frame percentage >50%, and in-frame site percentage ≥30%) and a full-ouORF approach (criteria: ≥15 aa, in-frame count ≥30, in-frame percentage >40%, and in-frame site percentage ≥30%). An ouORF was considered translated if it met either criterion (**Figs. S1 and S2**). nuORFs were identified using the 5′-UTR-only approach.

### Removal of uORFs resulting from isoform artifacts and consolidation of indistinguishable versions

To prevent the identification of uORFs that are part of the mORF in an alternative transcript isoform, nuORFs were removed if they overlapped another isoform’s mORF in any frame, and ouORFs were removed if they overlapped the mORF of another isoform in the same reading frame or if their start codon fell within the mORF of another isoform regardless of the frame. Isoform-specific versions of the same uORF, which shared a start or stop codon but differed because of alternative splicing, were then evaluated as follows: uORFs with indistinguishable P-site evidence were consolidated into a single representative, whereas those with distinct P-site evidence were retained separately.

### Identification of uORFs with CAGE-supported transcription start sites

CAGE-seq data from Arabidopsis Col-0 seedlings (56) (GEO accession GSE136356) were used to estimate the fraction of transcripts containing each uORF. For each uORF, we calculated %uORF+ as the fraction of CAGE signal originating upstream of the uORF start codon. The numerator was the total CAGE signal from 100 nt upstream of the annotated transcription start site (TSS) to the nucleotide immediately preceding the uORF start codon. The denominator was the total CAGE signal from the same 5′ boundary to the 3′ end of the mORF coding sequence. Thus, %uORF+ approximates the fraction of transcripts that initiate upstream of, and therefore contain, the uORF. uORFs with %uORF+ ≥ 30% were retained for downstream analyses.

### Determination of translation efficiency

Ribo-seq and RNA-seq abundances over the annotated CDS for each gene were quantified using RSEM (57). Translation efficiency (TE) was then calculated as the ratio of Ribo-seq to RNA-seq abundance and analyzed as log₂(TE + 0.01). A pseudo-count of 0.01 was added to avoid log₂(0), which is undefined.

### Kozak context and sequence logos

A reference position-specific base-frequency matrix was built from the -3 to +5 start-codon windows of the all genes with an annotated 5’ UTR; base counts at each position were added by a 0.01 pseudo-count, converted to per-position frequencies, and then divided by the most frequent base at that position, so each entry is a relative frequency ranging from 0 (least-preferred base) to 1 (most preferred base). For every start codon, the Kozak relative-strength score was computed as the geometric mean of these relative frequencies for the bases present at the five most informative flanking positions (−3, -2, -1, +4, +5), excluding the invariant AUG core; the score ranges from 0 to 1, and because it is a geometric mean a single strongly non-canonical position lowers it, so a high score requires all six positions to be favorable. Sequence logos over the -4 to +6 window (information content in bits, and base probability) were plotted with ggseqlogo (58).

### Gene ontology (GO) functional enrichment

Gene Ontology (GO) over-representation analysis was performed for three CAGE-supported uORF gene sets (genes containing ≥1 ouORF, ≥1 nuORF, or both) using all expressed genes with GO annotations as the background. The analysis was performed with topGO (59) using the complete Arabidopsis GO annotation for all three ontologies, the weight01 algorithm, and Fisher’s exact test (raw P < 0.01). The weight01 algorithm propagates significance through the GO graph while down-weighting parent terms explained by significant descendant terms. Significant GO terms were then collapsed into representative groups based on semantic similarity using rrvgo (60). The reduced GO results for the three uORF gene sets are provided in **Table S3**.

### Predicted protein-domain content and subcellular localization

To evaluate the consequences of ouORF-mediated N-terminal truncation, the full-length and N-terminally truncated mORF protein sequences (the truncated form beginning at the first in-frame AUG downstream of the ouORF; CAGE-filtered ouORFs only) were analyzed for both protein-domain content and predicted subcellular localization.

Pfam domain content was assessed with hmmscan (HMMER) (61) against a Pfam-A database (62)(https://ftp.ebi.ac.uk/pub/databases/Pfam, using Pfam-A.hmm.gz file from version 38.2), reporting only domains that met the Pfam curated gathering thresholds (--cut_ga). For each protein, the Pfam domain families detected in the full-length sequence were compared with those detected in the N-truncated sequence, and a domain was considered lost when its family was present in the full-length protein but no longer detected after truncation (**Table S4**).

Predicted subcellular localization was assessed with DeepLoc 2.1 (33) (https://services.healthtech.dtu.dk/services/DeepLoc-2.1/) using the high-sensitivity model and the short output format, applied to the same full-length and N-truncated sequences. For each protein, the localization predicted for the full-length protein was compared with that predicted for the N-truncated protein, and the number of proteins and their corresponding genes whose predicted localization changed after truncation was recorded (**Table S5**).

### Dual-luciferase reporter assay

The effect of ouORFs on mORF translation was assessed using a dual-luciferase reporter in pHsu133, a modified pGreenII 0800-LUC backbone containing *35S::Fluc* (*firefly luciferase*) and *35S::Rluc* (*Renilla luciferase*) (9). For each of the three ouORF-containing genes (*AT1G33250*, *AT3G61770*, and *AT5G20070*), 5′ UTR fragments containing either the wild-type or a mutated ouORF (start codon changed from ATG to AGG) and extending through the first four codons of the mORF were synthesized (Bio Basic) and cloned upstream of, and in frame with, the *Fluc* reporter via BamHI and NcoI. Fluc reported mORF translation, whereas Rluc served as an internal control. The synthesized sequences are provided in **File S1**.

The dual-luciferase assay was performed in protoplasts from *Arabidopsis thaliana* T87 suspension cell line (rpc00008, provided by the RIKEN BRC through the National BioResource Project of the MEXT, Japan). The cell lines were maintained in 50 ml liquid media (4.73 g/L LS (Caisson, LSP03-50LT), 3% sucrose, 0.00002% Thiamine (B1), 0.002% Myoinositol, 0.018% KH2PO4, and 0.2mg/L 2,4-D) in a 250 ml baffled Erlenmeyer flask shaken at 140 rpm at 22°C in the dark and subcultured every 7 days.

Protoplasts were isolated from cells 4 days after subculture (3 mL of a 7-day-old culture transferred to 50 mL of fresh medium). Cells were pelleted by centrifugation at 100 g for 3 minutes, resuspended, and digested in enzyme solution (1% [w/v] cellulase, 0.25% [w/v] macerozyme, 0.4 M mannitol, 20 mM KCl, 20 mM MES, 10 mM CaCl₂) for 2.5 h with gentle shaking (40 rpm) in the dark. Protoplasts were filtered through a 40 µm cell strainer, centrifuged at 100 × g at 4 °C, washed twice with cold W5 solution (154 mM NaCl, 125 mM CaCl₂, 5 mM KCl, 2 mM MES [pH 5.7], 5 mM glucose, 200 µM KNO₃), counted with a hemocytometer, and resuspended in cold MMG solution (4.5 mM MES [pH 5.7], 0.4 M mannitol, 15 mM MgCl₂) at 10⁵ protoplasts per 150 µL.

Transformation and the luciferase assay were performed as described by (63). For each of 6–12 biological replicates, 10⁵ protoplasts were mixed with 5 µg of plasmid DNA (prepared with a Zymo midiprep kit [Zymo D4201]) and 170 µL of PEG solution (40% [w/v] PEG4000, 0.2 M mannitol, 100 mM CaCl₂) and incubated for 5 min. After washing and overnight incubation in the dark, each transformed sample was split into two equal aliquots: one for the luciferase assay and one for RNA extraction. Luciferase activities were measured with the Dual-Luciferase Reporter Assay System (Promega, E1960) on a GloMax Navigator plate reader (Promega, GM2010) according to the manufacturer’s protocol. Fluc relative luminescence is reported as Fluc luminescence normalized to Rluc luminescence for each sample.

### RNA extraction and RT-qPCR

To assess whether the differences in reporter activity reflected translational rather than transcript level changes, reporter mRNA levels were quantified by RT-qPCR. For each construct, the RNA aliquots from two transformation samples were combined, and total RNA was extracted by the TRIzol method and purified with the RNA Clean & Concentrator-25 kit (Zymo R1018). RNA was treated with 6U of TURBO DNase (Invitrogen AM2238) at 37 °C for 30 min to remove DNA and repurified using the RNA Clean & Concentrator-5 kit (Zymo R1016). cDNA was synthesized using the LunaScript RT SuperMix Kit (NEB, E3010) in a 15 µL reaction. Twofold-diluted cDNA was used as the template for RT-qPCR with Luna Universal qPCR Master Mix (NEB, M3003) in a final reaction volume of 10 µL with three technical replicates. Relative *Fluc* expression was calculated using the ΔΔCt method with *Rluc* as the reference gene. Primer pairs were as follows: Fluc, 5′-AAGTACTCAGCGTAAGTGAT-3′ and 5′- CCATGGAAGACGCCAAAAACAT-3′; Rluc, 5′-CATGGGATGAATGGCCTGATATTG-3′ and 5′- GATAATGTTGGACGACGAACTTC-3′.

### In vitro transcription, translation, and immunoblotting

To test the effect of ouORFs on the production of N-terminally truncated proteins, capped and polyadenylated mRNAs were synthesized in vitro for *SCR* (AT3G54220), *mTERF3* (AT2G36000), and *ATPS2* (AT1G19920). For each gene, DNA fragments containing a T7 promoter, a 5′ UTR containing either the wild-type or a mutated ouORF (start codon changed from ATG to AGG), and the first 93–171 codons of the mORF fused to a C-terminal HA tag were synthesized (Bio Basic) (**File S2**). The fragment length was chosen to ensure that the N-terminal truncation would result in proteins of at least 5 kDa for reliable detection. DNA fragments were PCR-amplified, purified using the DNA Clean & Concentrator-5 Kit (Zymo, D4013), and 1 µg of purified PCR product was used as the template for in vitro transcription.

In vitro transcription was performed using the HiScribe T7 High Yield RNA Synthesis Kit (NEB, E2040S) at 37°C for 2 h. The synthesized RNA was treated with 5 U DNase I (Zymo, E1010) at room temperature for 15 min and purified using the RNA Clean & Concentrator-25 Kit (Zymo, R1018). Approximately 25 µg of RNA was then capped using the ScriptCap m⁷G Capping System (Cellscript, C-SCCE0625; 2 µL capping enzyme per reaction) at 37°C for 30 min and polyadenylated using the A-Plus Poly(A) Polymerase Tailing Kit (50 U, Cellscript, C- PAP5104H) at 37°C for 30 min. The RNA was purified again using the RNA Clean & Concentrator-25 Kit. Approximately 500 ng of capped and polyadenylated mRNA was translated in vitro using wheat germ extract (Promega, L4380) in a 20 µL reaction at 25°C for 2 h.

In vitro transcription was carried out using the HiScribe T7 High Yield RNA Synthesis Kit (NEB, E2040S) at 37°C for 2 hours. The synthesized RNA was treated with 5 U of DNase I (Zymo E1010) at room temperature for 15 min and purified using the RNA Clean & Concentrator-25 kit (Zymo R1018). Subsequently, approximately 25 µg of RNA was capped with the ScriptCap m⁷G Capping System (CELLSCRIPT, C-SCCE0625; 2 µL capping enzyme per reaction) at 37°C for 30 min and polyadenylated with the A-Plus Poly(A) Polymerase Tailing Kit (50U, CELLSCRIPT, C-PAP5104H) at 37 °C for 30 min. The RNA was then purified with the RNA Clean & Concentrator-25 kit. Approximately 500 ng of mRNA was translated in vitro using wheat germ extract (Promega, L4380) containing 50 mM potassium acetate in a final reaction volume of 20 µL at 25°C for 2 h.

Translation products (10 µL per sample) were separated on 16% (w/v) Tris–Tricine PAGE gels (Invitrogen, EC66952BOX) and transferred to 0.2 µm nitrocellulose membranes at 30 V for 90 min in Tris-glycine transfer buffer containing 20% (v/v) methanol. Membranes were blocked with 5% (w/v) non-fat milk in TBST (TBS containing 0.05% Tween-20) for 1 h at room temperature with gentle shaking and incubated overnight at 4°C with an anti-HA primary antibody (rat monoclonal, clone 3F10; Roche, 11867423001) diluted 1:2,000. After washing, membranes were incubated with an HRP-conjugated goat anti-rat IgG (H+L) secondary antibody (Agrisera, AS10 1187) diluted 1:10,000 for 30 min at room temperature with gentle shaking. Signals were detected using Amersham ECL Prime Western Blotting Detection Reagent (Cytiva, RPN2232) and imaged on an Azure Biosystems imager in cumulative mode. Band intensities were quantified as peak areas in ImageJ (64), and the relative abundance of the full-length and N-terminally truncated isoforms was presented as the percentage of the total signal (full-length + truncated) to compare between wild-type and mutant ouORF constructs.

### Subcellular localization and confocal microscopy

Four ATPS2–GFP constructs were generated: WT ouORF (the 5′ UTR containing the ouORF and extending through the first 51 amino acids of the mORF), mutated ouORF (the same 5′ UTR with the ouORF start codon changed from ATG to TAG), Δ5′ UTR (lacking the ATPS2 5′ UTR), and ΔN51 (lacking both the 5′ UTR and the first 51 amino acids of the mORF). The corresponding DNA fragments (**File S3**) were synthesized by Bio Basic and cloned upstream of GFP in the 35S-sfGFP-nosT vector (Addgene plasmid #80129; Fujii and Kodama 2015) using the SalI and NcoI restriction sites. All plasmids were prepared using a ZymoPURE II Plasmid Midiprep Kit (Zymo, D4201).

Mesophyll protoplasts were isolated from 4-week-old *Arabidopsis thaliana* Col-0 plants grown in soil under a 12 h light / 12 h dark photoperiod at 100 µmol m⁻² s⁻¹. The protoplasts were isolated as described (63), with additional purification by Percoll density gradient centrifugation following the protocol of (65). For each transformation, 15 µg of plasmid DNA was transformed into 200 µL of protoplast suspension (1 × 10⁶ cells/mL) using the PEG method with an 8 min incubation. Transformed protoplasts were incubated overnight in a Percival growth chamber and imaged on a Zeiss LSM 980 confocal microscope equipped with Airyscan using 488 nm excitation and emission ranges of 489–541 nm for GFP and 668–758 nm for chlorophyll autofluorescence. For each construct, GFP localization in individual cells was classified as chloroplast-only, cytosol-only, or dual localization. A total of 50–100 cells per construct were scored across five independent experiments performed on different days.

### AI usage

The authors used large language models (Claude, ChatGPT, Gemini, and Grok) to assist with manuscript editing for grammar and clarity and to improve analysis code according to authors’ instructions. These tools were not used to generate data or draw scientific conclusions. All outputs were reviewed and verified by the authors, who take full responsibility for the content of the manuscript.

## AUTHOR CONTRIBUTIONS

HLW and PYH conceived the research. HLW developed methods for ouORF identification, performed the bioinformatics analyses, and prepared the figures and tables. YC performed the dual-luciferase assays, RT-qPCR, immunoblotting, and subcellular localization assays. IK performed the initial subcellular localization assays. QA performed the initial dual-luciferase assays. PYH performed the initial immunoblotting. HLW and PYH wrote the manuscript with input from all authors.

## ACKNOWLEDGEMENT

We thank Hideki Takahashi for the helpful discussion on the research. This work used computational resources and services provided by the Institute for Cyber-Enabled Research at Michigan State University. This work was supported by a predoctoral training award under Grant Number T32-GM110523 from the National Institute of General Medical Sciences of the National Institutes of Health to IDK, and research grants from the National Science Foundation under Award Numbers 2425390 and 2051885, and the National Institute of General Medical Sciences of the National Institutes of Health under Award Number R35GM155375 to PYH. The content is solely the responsibility of the authors and does not necessarily represent the official views of the NSF or NIH.

---

**Fig. S1.**
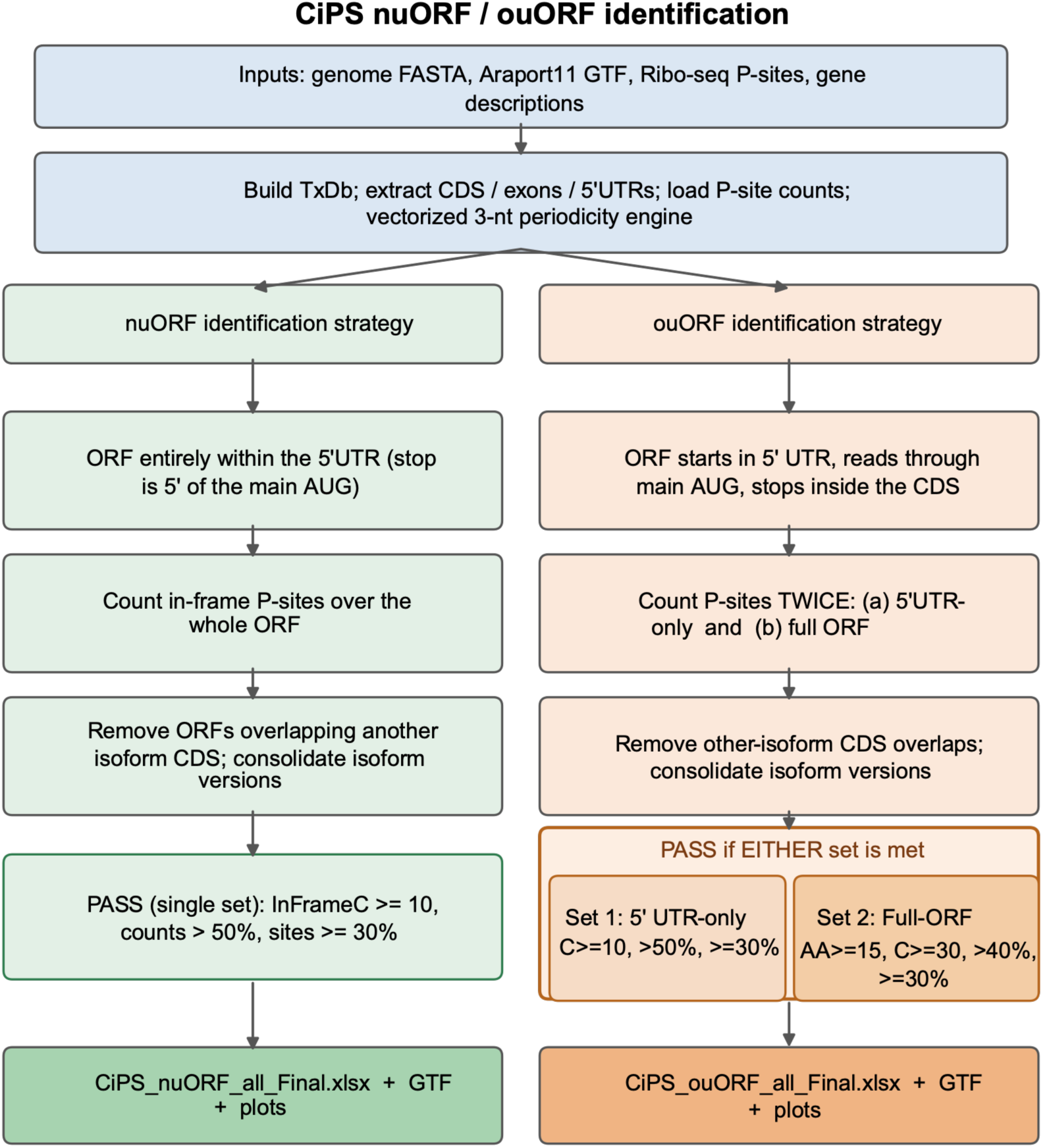
CiPS nuORF/ouORF identification workflow. The workflow begins with building transcript models, 5’ UTR annotations, and ribosome P-site positions, then uses a vectorized three-nucleotide periodicity engine to score candidate ORFs. nuORFs and ouORFs are processed in parallel. ouORFs are evaluated using both the 5’ UTR-only and whole-ouORF approaches to capture those with ribosome density biased between the 5’ UTR and the mORF. The pipeline then removes isoform artifacts, consolidates indistinguishable uORFs, and exports nuORF/ouORF tables, GTF annotations, and diagnostic plots.

**Fig. S2.**
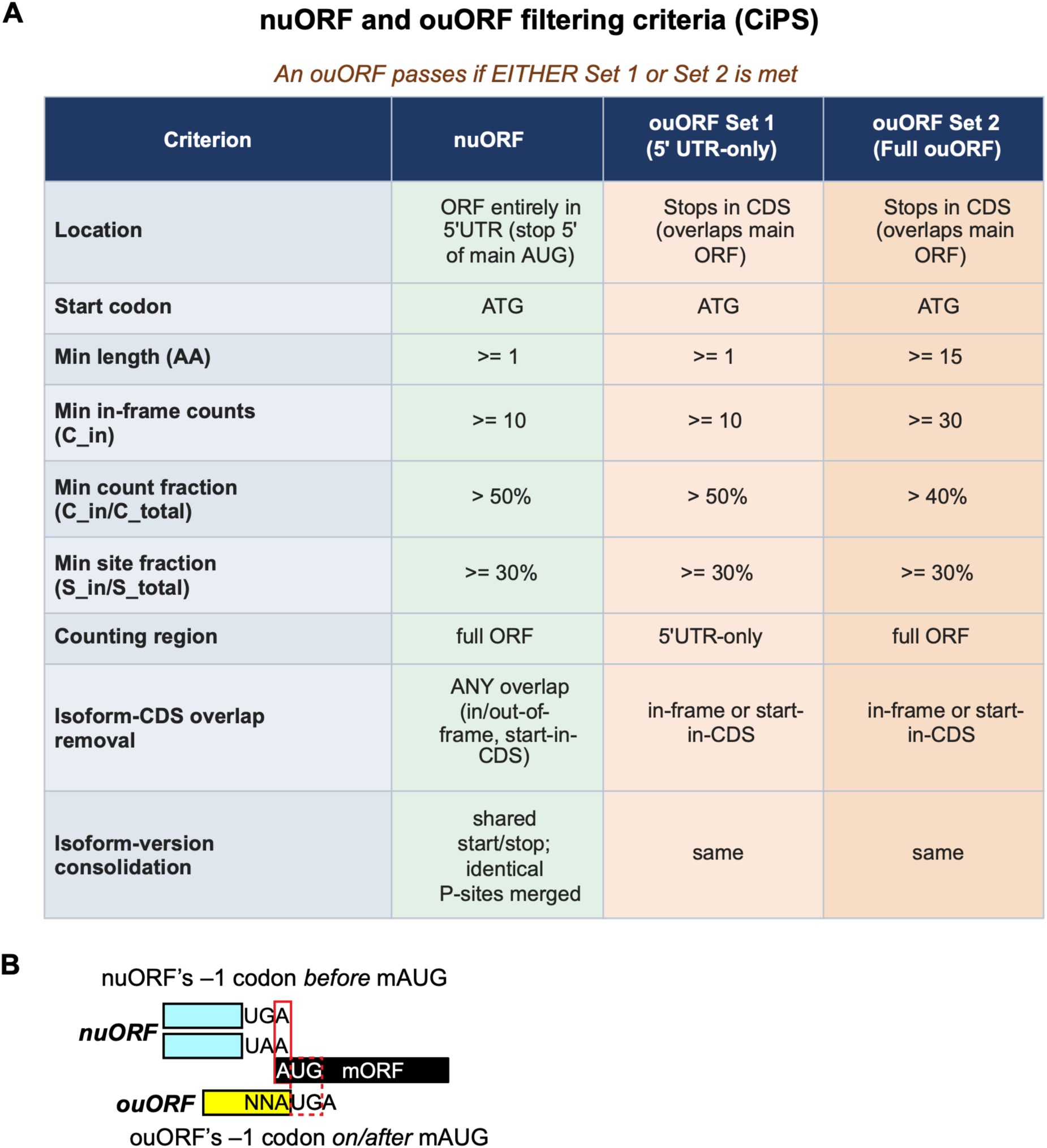
Criteria for nuORF and ouORF identification. (A) Detailed criteria used for nuORF and ouORF identification with CiPS. (B) Classification of nuORFs and ouORFs located near the mORF start codon.

**Fig. S3.**
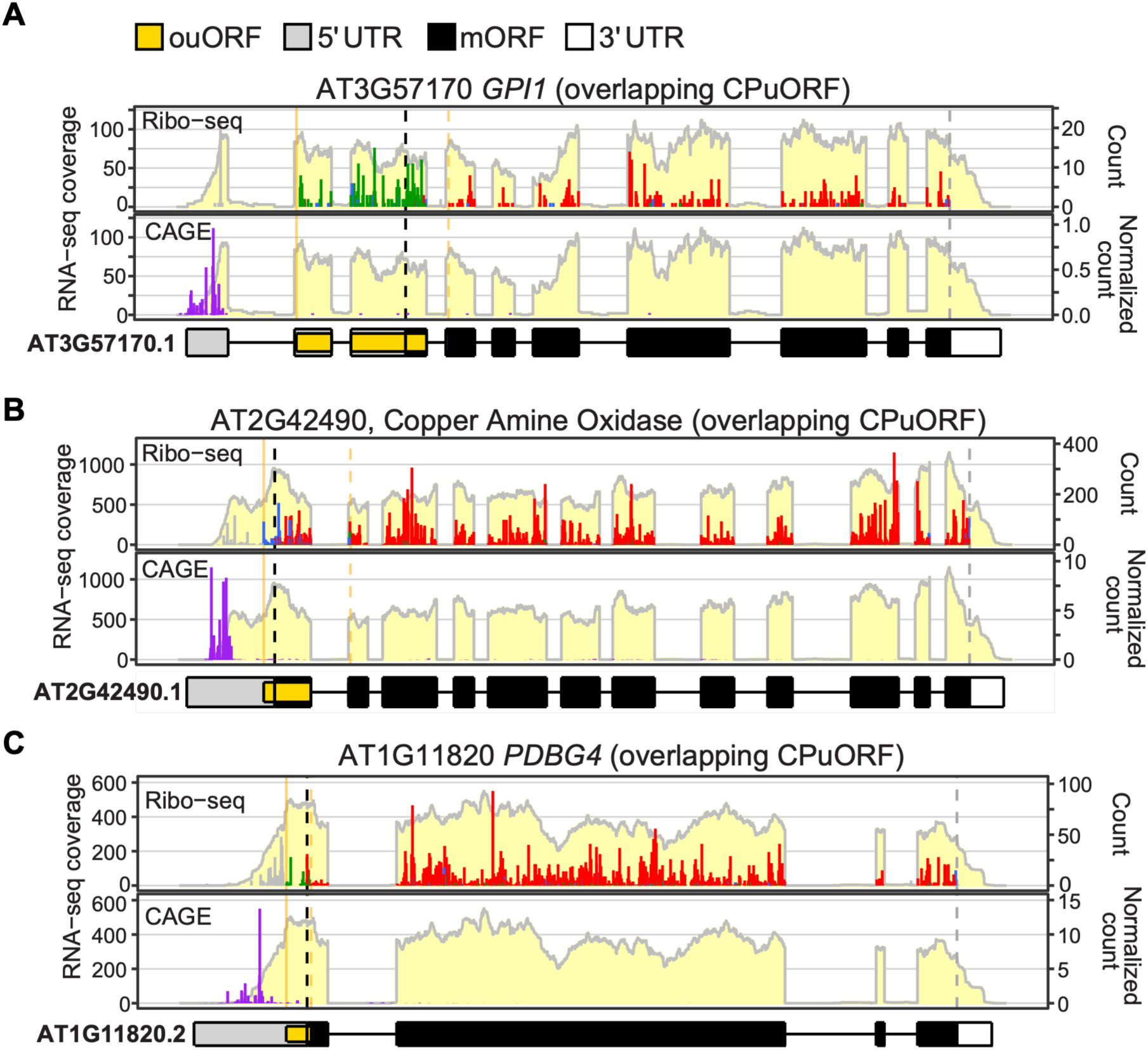
Expression profiles of three conserved peptide overlapping uORFs (CPouORFs). RNA-seq, Ribo-seq, and CAGE-seq profiles of three CPouORF-containing genes visualized using *ggRibo* (66). S3A top panel is also shown in Figure 1G. The three CPouORFs from A to C have 97.7, 96,5, 97.3% ouORF+, respectively. Data presentation is described in the Fig. 1E legend.

**Fig. S4.**
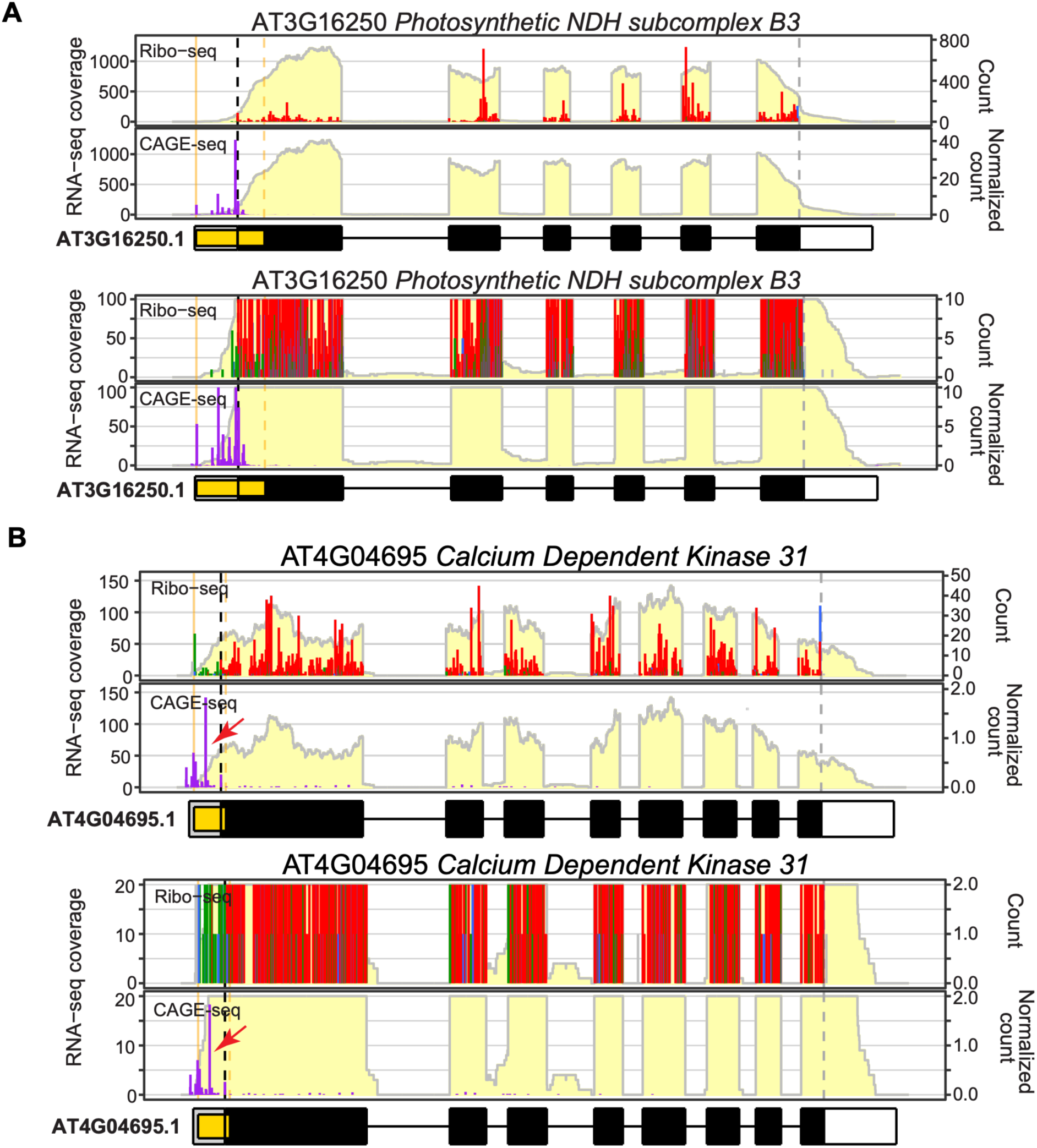
Transcription start site selection strongly influences ouORF inclusion. (A, B) Examples of ouORF-containing genes illustrating the effect of transcription start site selection on ouORF inclusion. RNA-seq, Ribo-seq, and CAGE-seq data are visualized using *ggRibo*. The lower plots in each panel show the same data as the upper plots but with a zoomed-in view at the y-axis. Data presentation is described in the Fig. 1E legend.

**Fig. S5.**
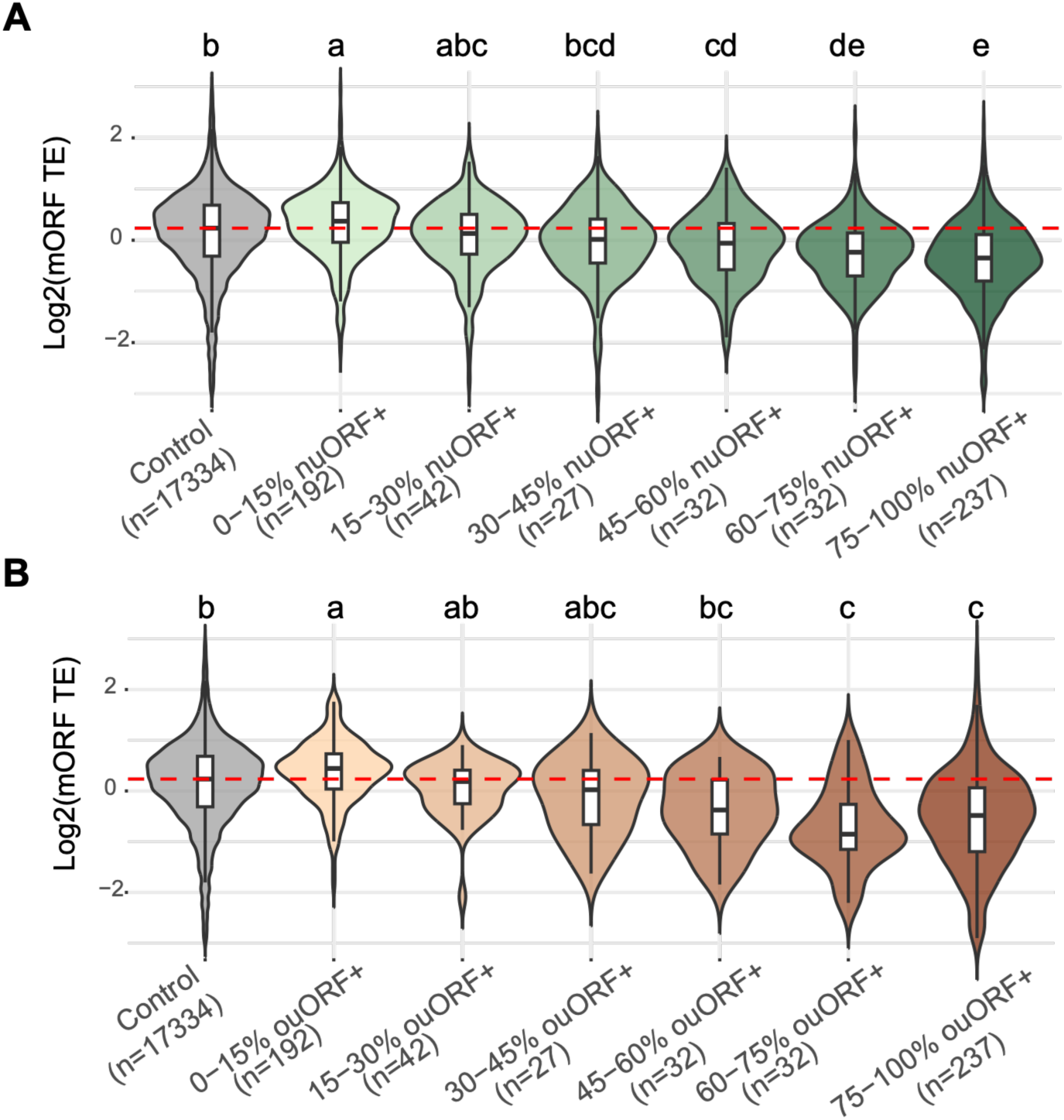
TSS-mediated uORF inclusion is tightly associated with translational repression. Distribution of mORF translation efficiency (TE) for genes lacking uORFs and genes with different percentages of nuORF-containing (A) or ouORF-containing (B) transcripts, based on the proportion of CAGE signal upstream of the corresponding uORF start codon. Repression increases when TSS support increases, indicating that uORFs are most repressive when retained in a larger fraction of expressed transcripts. Panel B is identical to Fig. 2A, but with gene numbers shown for each group. The red dashed line indicates the median TE of genes lacking uORFs. Different letters indicate statistically significant differences (ANOVA followed by Tukey’s HSD test, p < 0.05).

**Fig. S6.**
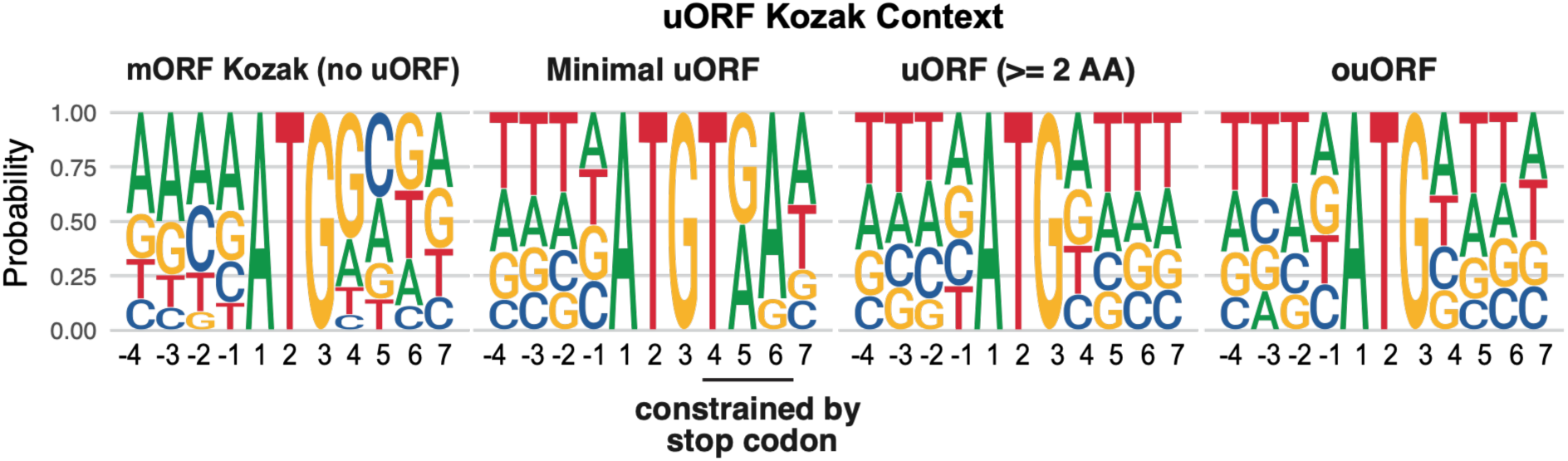
uORFs exhibit weaker Kozak contexts than mORFs. Sequence logos surrounding the start codons of mORFs from uORF-less genes, minimal nuORFs, nuORFs ≥2 aa, and ouORFs. The data are identical to those in Fig. 4A, but the logos display nucleotide probabilities instead of information content (bits).

**Fig. S7.**
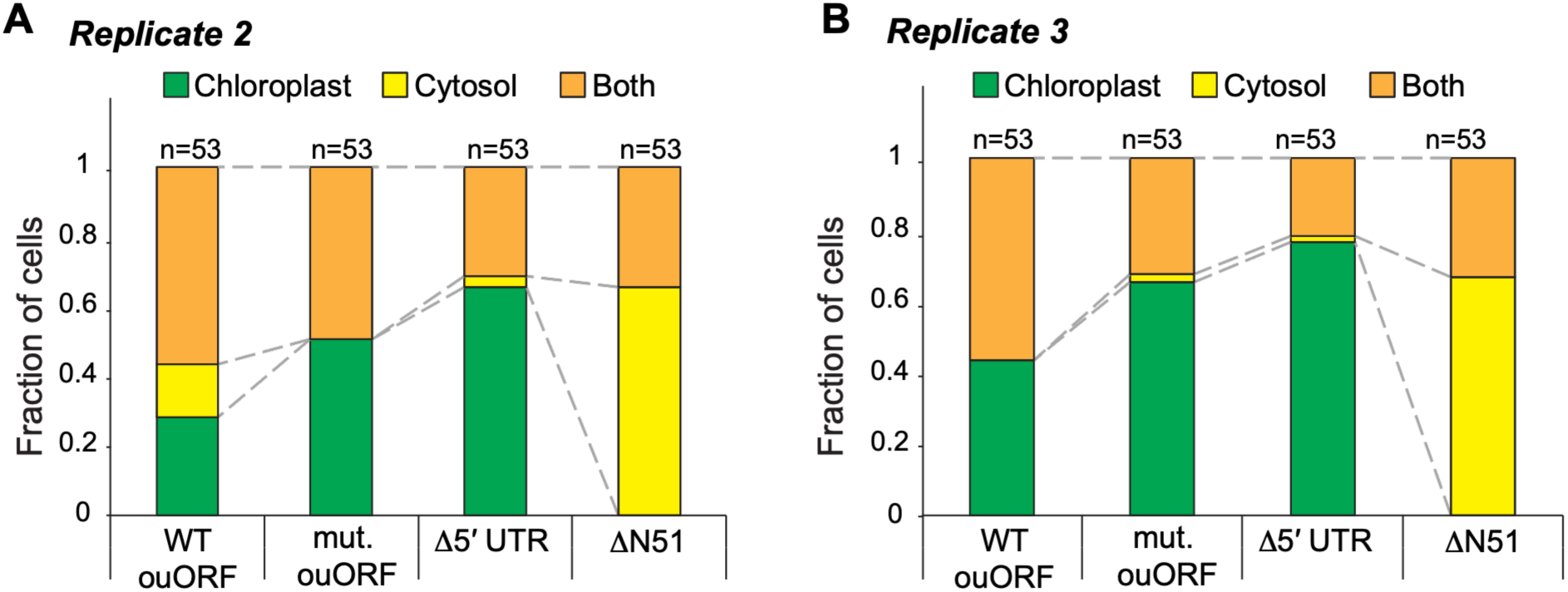
Independent quantification of ATPS2–GFP subcellular localization. (A, B) Additional independent replicate quantifications of ATPS2–GFP localization in Arabidopsis leaf protoplasts expressing the wild-type ouORF, mutated ouORF, Δ5′ UTR, or ΔN-terminal constructs. Cells were classified as chloroplast-only, cytosol-only, or dual-localized. Numbers above the bars indicate the number of cells analyzed for each construct.

## SUPPLEMENTAL TABLES

**Table S1. Translated ouORFs (mORF-overlapping uORFs) detected by CiPS.** (Table_S1_CiPS_ouORF_Final.xlsx).

**Table S2. Translated nuORFs (non-overlapping uORFs) detected by CiPS.** (Table_S2_CiPS_nuORF_Final.xlsx). Columns are described in Table S1.

**Table S3. Gene Ontology over-representation for the CAGE-supported uORF gene sets.** (Table_S3_GO_uORF.xlsx).

**Table S4.** Predicted ouORF mediated domain loss (Table_S4_ouORF_domain_loss.xlsx).

**Table S5.** Predicted ouORF mediated localization change (Table_S5_ouORF_DeepLoc_localization_change.xlsx).

